# A CRISPR-CAS9 high throughput machine-learning platform for modulation of genes involved in Parkinson’s disease-associated PINK1-mitophagy in iPSC-derived dopaminergic neurons

**DOI:** 10.1101/2025.06.10.658840

**Authors:** Marc P. M. Soutar, Benedetta Carbone, Petya Kindalova, Rahil Mehrizi, Fernanda Martins Lopes, Nathaniel Lam, Alice Rockliffe, Toni Braybrook, Matthew Taylor, Cuong Nguyen, Fiona Ducotterd, Alastair D. Reith, Lisa Mohamet, Helene Plun-Favreau

**Affiliations:** Queen Square Institute of Neurology, Department of Neurodegenerative Disease, University College London, London, WC1N 3BG, United Kingdom; GlaxoSmithKline R&D, Gunnels Wood Rd, Hertfordshire, Stevenage SG1 2NY UK; ARUK Drug Discovery Institute, The Cruciform Building Department of Neuromuscular Disorders University College London, Gower Street, London, WC1E 6BT, United Kingdom; Breckenfield Consulting Ltd, London, UK

**Author notes:** contributed equally (Lisa Mohamet, Helene Plun-Favreau). Co-first author (Marc P. M. Soutar, Benedetta Carbone). **<u>Competing Interest Statement</u>** BC, PK, RM, FML, NL, AR, TB, MT, CN, ADR, LM are or were employees of GSK, a global healthcare company that may conceivably benefit financially through this publication.

## Abstract

Parkinson’s disease (PD) is a progressive neurodegenerative disorder characterised by the loss of dopaminergic neurons, driven by complex molecular mechanisms that are not fully understood. To address this issue, we have developed a novel high-content phenotypic screening platform using human induced pluripotent stem cell-derived dopaminergic neurons to investigate the PINK1-PARKIN mitophagy pathway, a critical process in PD pathogenesis.

Utilising high throughput, 384 well arrayed CRISPR-CAS9 genetic manipulation and high-content immunofluorescence imaging complemented with machine learning analysis, we examined ubiquitin (Ub) pSer65 levels. Ub pSer65, a potential PD clinical biomarker, is a key marker of mitophagy initiation in dopaminergic neurons upon mitophagy initiation using exogenous stimuli to mimic the disease relevant environment. The CRISPR-CAS9 knockout (KO) screen revealed two distinct phenotypic classes: essential genes causing cell death upon deletion, and genes modulating Ub pSer65 levels. Notably, KO of *PINK1, PARKIN*, and *TOM7 genes* decreased Ub pSer65 upregulation during mitophagy activation, confirming their established roles in the pathway and validating the suitability of the platform for target identification.

This innovative platform provides a precise tool to further interrogate PD-associated genes, offering insights into mitophagy-related pathogenic mechanisms and identification of potential therapeutic targets. By bridging functional genomics with disease-specific neuronal models, this approach presents a promising strategy for advancing PD research and developing targeted interventions. To our knowledge, this is the first reported use of a human, translationally relevant cell model to study genetic perturbation within a disease relevant phenotype.

## Introduction

Parkinson’s disease (PD) is a neurodegenerative disorder characterised by the progressive loss of dopaminergic (DA) neurons in the substantia nigra, leading to motor and non-motor dysfunction, as well as cognitive impairment (Gonzalez-Casacuberta, Juarez-Flores et al. 2019). Despite extensive research, the molecular mechanisms underlying PD pathogenesis remain incompletely understood, hindering the development of targeted therapeutic interventions with significant disease modifying efficacy.

Clinical genetics plays a crucial role in understanding PD pathogenesis, with rare Mendelian genes revealing key cellular pathways (Gonzalez-Casacuberta, Juarez-Flores et al. 2019). Notably, mutations in autosomal recessive Mendelian genes, such as PTEN-induced putative kinase 1 (*PINK1*) and *PARKIN*, encoding a Ser/Thr protein kinase, and an E3 ubiquitin ligase have been linked to defects in the selective degradation of damaged mitochondria through autophagy (mitophagy) (Kitada, Asakawa et al. 1998, Valente, Abou-Sleiman et al. 2004, Valente, Salvi et al. 2004, Tsukada, Kanazawa et al. 2016). PINK1 and PARKIN are essential in initiating mitophagy, ensuring the clearance of heavily dysfunctional mitochondria while maintaining a healthy mitochondrial network) (Jin, Lazarou et al. 2010, Narendra, Jin et al. 2010, Narendra and Youle 2024).

When mitochondria are damaged, PINK1 accumulates on the outer mitochondrial membrane (Narendra, Jin et al. 2010) where it phosphorylates ubiquitin at Ser65 (Ub pSer65). This modification serves as a receptor for Parkin recruitment from the cytosol to the damaged mitochondria, where Parkin is subsequently phosphorylated at Ser65 in a PINK1-dependent manner, activating its E3 ubiquitin ligase activity (Kazlauskaite, Kondapalli et al. 2014, Shiba-Fukushima, Arano et al. 2014, Fiesel, Ando et al. 2015, Fiesel and Springer 2015, Lazarou, Sliter et al. 2015, Watzlawik, Hou et al. 2021). Activated Parkin ubiquitinates a wide range of outer mitochondrial proteins, marking the damaged mitochondria with Ub pSer65 chains, and leading to their lysosomal degradation (Narendra, Tanaka et al. 2008, Narendra, Jin et al. 2010, McWilliams and Muqit 2017). Mutations in PINK1 and PARKIN can impair mitophagy, leading to the accumulation of damaged mitochondria (Matsuda, Sato et al. 2010). Elevated levels of Ub pSer65 have been observed in post-mortem brain samples, as well as in human cells and blood samples from PD patients, with a correlation to disease progression (Fiesel, Ando et al. 2015, Shiba-Fukushima, Ishikawa et al. 2017, Watzlawik, Hou et al. 2021, Hertz, Chin et al. 2024). Moreover, DA neurons from sporadic PD patients exhibit a significant reduction in PINK1 and PARKIN protein compared to controls (Chen, McDonald et al. 2023). Mitochondrial toxins have been shown to induce Parkinsonism in both human and animal models (Tsukada, Kanazawa et al. 2016), strengthening the connection between mitophagy dysfunction and PD. Collectively, these findings underscore the role of PINK1-mediated mitophagy in PD development and highlight the potential of Ub pSer65 as a clinical biomarker (Fiesel and Springer 2015). Thus, enhancing effectiveness of the mitophagy pathway offers a promising strategy for therapeutic intervention in PD (Miller and Muqit 2019, Aman, Ryan et al. 2020, Clark, Vazquez de la Torre et al. 2021, Malpartida, Williamson et al. 2021).

While the identification of novel modulators for development of mitophagy-targeted therapeutics is challenging, recent research suggests that drug targets that are supported by human genetic evidence of disease association have a two-fold increased likelihood of a successful clinical outcome (Nelson, Tipney et al. 2015, Minikel, Painter et al. 2024). PD genome-wide association studies (GWAS) studies have identified several risk loci (Leonard 2025). However, translating these findings into new treatments remains a considerable challenge, necessitating the development of advanced functional genomic technologies in disease-relevant cell models.

In recent high-content screening studies, Ub pSer65 has been employed as an output to evaluate potential compound activators of PARKIN in fibroblasts and SHSY5Y cells (Fiesel, Ando et al. 2015, Fiesel and Springer 2015, Giorgianni and Beranova-Giorgianni 2016, Shiba-Fukushima, Ishikawa et al. 2017, Ordureau, Paulo et al. 2018, Watzlawik, Hou et al. 2021, Chen, McDonald et al. 2023, Tufi, Clark et al. 2023). In this study, we developed and validated a scalable, high-content phenotypic arrayed screening *in vitro* platform that uses human induced pluripotent stem cells (hiPSC) to generate dopaminergic neurons (iPSC-DA neurons), the key disease-relevant key cell type responsible for motor dysfunction in PD. This platform is designed for high-throughput screening and genetic manipulation via CRISPR-CAS9, allowing for systematic exploration of genes modulating PINK1-dependent Ub pSer65 accumulation (induced by oligomycin/antimycin (O/A) combination), a marker of mitophagy initiation. Oligomycin (an ATP synthase inhibitor) and antimycin (a complex III inhibitor) are well-established inducers of mitochondrial stress, which trigger mitophagy activation (Soutar, Kempthorne et al. 2019).

For validation of this cell assay platform, we selected a panel of 12 targets: PINK1, PARKIN, TOM7, SYNJ2 and SYNJ2B, USP30, KANSL1, KAT8, OGT, ULK1, EIF2AK1 and FBOX7, based on their reported regulation of Ub pSer65, from which PINK1, PARKIN and USP30’s role in Ub pSer65 regulation has been demonstrated in DA neurons (Burchell, Nelson et al. 2013, Shiba-Fukushima, Ishikawa et al. 2017, Ordureau, Paulo et al. 2018, Sekine, Wang et al. 2019, Hung, Lombardo et al. 2021, Tsefou, Walker et al. 2021, Harbauer, Hees et al. 2022, Maruszczak, Jung et al. 2022, Soutar, Melandri et al. 2022, Singh, Agarwal et al. 2024).

We investigated their role in our activated mitophagy platform by assessing Ub pSer65 modulation after KO using high-content immunofluorescence imaging and machine learning analysis.

CRISPR-CAS9 KO screen revealed two distinct phenotypic classes. The first class consisted of essential genes, such as *KANSL1*, KO of which induced cell death. The second class included Ub pSer65 modulators, where KO of *PINK1, PARKIN*, and *TOM7* prevented Ub pSer65 upregulation on mitophagy activation, confirming their previously characterised roles. In contrast, the KO of *USP30, SYNJ2*, and *SYNJ2BP* genes did not significantly affect Ub pSer65 levels compared to controls. These findings demonstrate the utility of the platform for evaluating candidate PD genes involved in mitophagy-related pathogenic mechanisms. It provides a precise tool for investigating genes associated with PD, offering insights into new therapeutic targets, mechanistic pathways, and downstream interventions with a higher likelihood of achieving clinical success.

## Results

The schematic in Fig. 1A summarises iPSC-DA neurons differentiation workflow adapted from (Nolbrant, Heuer et al. 2017) and describes the patterned expression of the floor plate midbrain markers in our primary iPSC donor line. The reference protocol establishes some criteria for assessment of a successful *in vitro* differentiation based on the human embryonic neural development. LMX1 expression indicates correct patterning towards the floor plate, whereas OTX2 and FOXA2 expression is localised to the forebrain-midbrain and midbrain-hindbrain regions, respectively. Only the simultaneous expression of all 3 markers indicates a successful midbrain, floor plate identity.

**Fig. 1:**
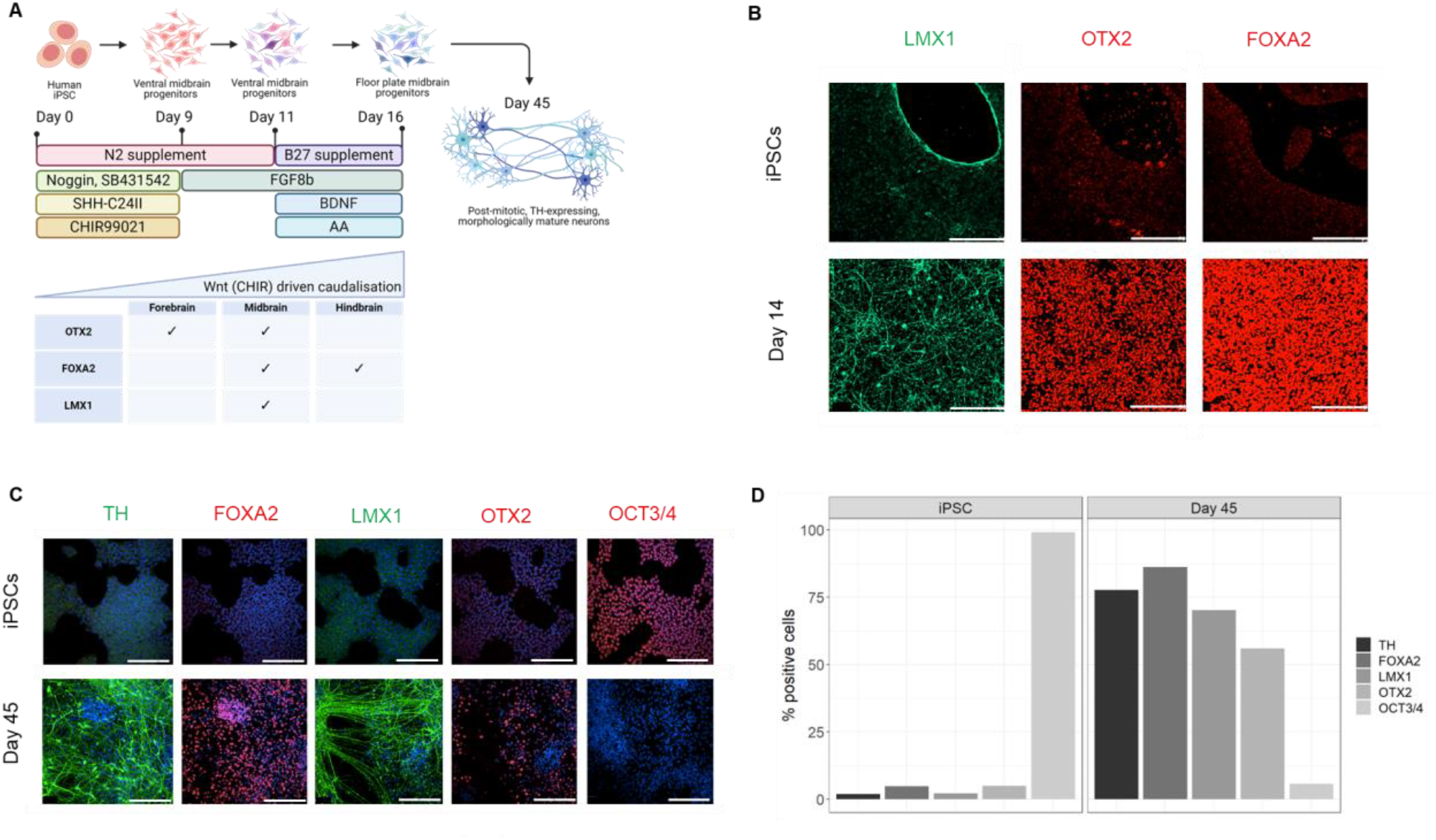
Phenotypic characterisation of iPSC DA neurons exhibit floor plate progenitor markers and mature TH expression on day 45. A) Schematic of the differentiation protocol from human iPSCs to DA neurons. Key stages, supplements, and regional marker expression patterns are shown. Created in BioRender. Carbone, B. (2025) https://BioRender.com/yupzpxi B) Immunofluorescence images of day 14 floor plate progenitor cells expressing FOXA2, LMX1, and OTX2 markers. Scale bars represent 200μm. C) Immunofluorescence images showing expression of TH, FOXA2, LMX1, OTX2, and OCT3/4 in iPSCs and day 45 neurons. Scale bars represent 200μm. D) Quantification of marker-positive cells at iPSC and day 45 neurons. The mean percentage of positive cells for each marker is shown, with 2 technical replicate wells available for each marker by experimental group combination.

The differentiation protocol employed dual SMAD inhibition, using Noggin and SB-431542 to promote neuronal induction (Nolbrant, Heuer et al. 2017). The addition of CHIR99021, a WNT pathway agonist, controls rostro-caudal differentiation in a gradient-dependent manner, whereas Sonic Hedgehog (SHH) directs ventral specification allowing floor plate patterning. The optimal concentrations of SHH and CHIR99021 capable of inducing an efficient DA neuron phenotype were experimentally determined in a donor-specific titration, and each donor line was assessed for differentiation/patterning efficiency (the data for differentiation optimisation of our primary donor STBCi101-A can be found in Supp. Fig. 1A). While the original study employed human embryonic stem cells (hESC) (Nolbrant, Heuer et al. 2017), we successfully adapted this for hiPSCs, leveraging the same optimisation techniques.

Donor-optimised DA neuron differentiation was characterised through immunocytochemical (ICC) analysis of key transcription factors associated with floor plate midbrain FOXA2, OTX2 and LMX1 in day 14 progenitor cells. As shown in Fig. 1B, day 14 progenitor cells from donor line STBCi101-A showed uniform expression of all three markers, significantly upregulated compared to undifferentiated iPSCs. Two additional healthy donors were differentiated towards DA neuron fate (Supp. Fig. 1B and C). We established a highly scalable and robust platform that enabled generation of >1billion neuronal progenitors (day 16) from a starting culture of 20 million undifferentiated iPSCs. IPSC-DA progenitors (D16) were subsequently differentiated until day 45, when they were assessed for the mature dopaminergic identity neuron marker tyrosine hydroxylase (TH), as well as the continued expression of FOXA2, OTX2 and LMX1 (Fig. 1C) The expression of these markers was quantified and compared against any non-differentiated iPSC control (Fig. 1D). OCT3/4 was used as a control marker for iPSC pluripotency, where its expression was completely absent in terminally differentiated day 45 neurons.

These data confirmed the reproducibility and robustness of the adapted protocol from hESCs to donor-derived hiPSC-DA neurons and established a large-scale *in vitro* platform for studying molecular mechanisms in patient-derived DA neurons *in vitro*.

Day 45 iPSC-DA neurons were subjected to a range of concentrations of O/A (0.0625 μM – 4 μM of each compound in a single treatment) for 5 hours to establish a robust assay window and response to treatment was detected via ICC through high-content confocal imaging (Fig. 2A).

**Fig. 2:**
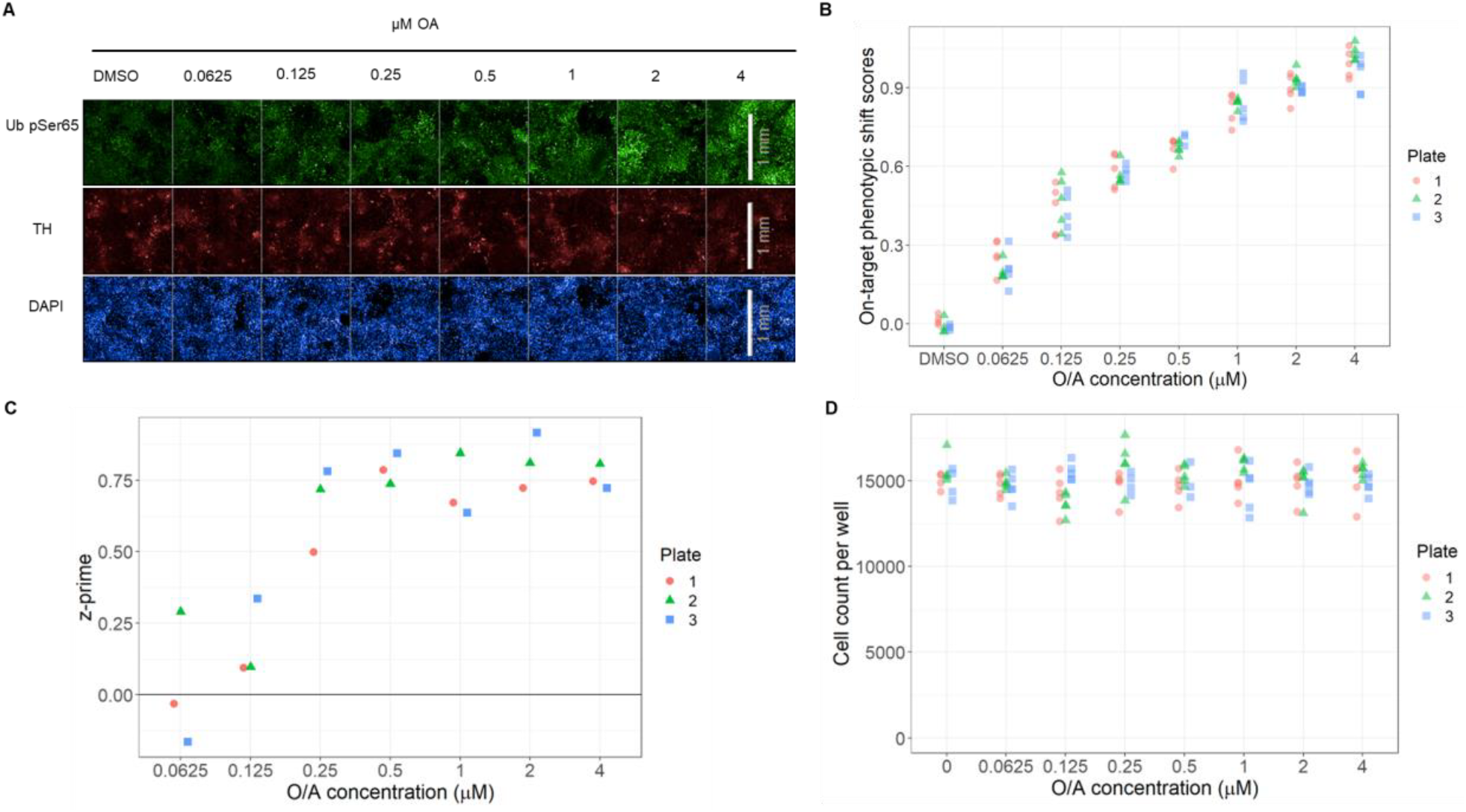
Assessment of activated mitophagy (Ub pSer65) in day 45 DANs - assay window across technical replicate plates. A) Representative immunofluorescence images showing Ub pSer65 signal in response to increasing concentrations of O/A treatment. Green: Ub pSer65, Red: TH, Blue: DAPI. Scale bars represent 1mm. B) Quantification of Ub pSer65 signal (on-target phenotypic shift) in response to O/A treatment across three replicate plates, as determined by AIML analysis of high-content imaging data. C) Z’ values calculated for each O/A concentration across the three technical replicate plates. The 0.5 μM dose (used in pilot) showed Z’ > 0.6 for all replicates. D) Cell counts per well across O/A concentrations, demonstrating no significant toxicity within the tested dose range and treatment duration. B-D) Same wells utilised for all three visualisations, with 3 to 5 replicates used per treatment by plate combination. Three wells were excluded from plate 3 due to cell count lower than 1000 cells per well (technical quality cut off), those wells were excluded from visualisations or Z’ calculation.

To fully capture this multidimensional data, we employed artificial intelligence/machine learning (AIML) approaches to collect maximum information content without the need for explicit biased segmentation rules (Mehrizi, Mehrjou et al. 2023). By doing so, treatment-induced changes could be quantified in cellular phenotypes in an unbiased manner. Briefly, a machine learning model was trained on acquired images using cell-centred patches from TH positive cells to produce a latent representation of the cell images. The model was trained to detect differences in Ub pSer65 between the basal levels of DMSO-treated cells and the levels present in O/A treated wells at maximum dose. The differences were captured as a series of imaging features such as signal intensity, area and granularity, among others, and were collectively referred to as the phenotypic shift (see Methods section for more details). The phenotypic shift was calculated among two perpendicular axis, with the on-target phenotypic shift representing changes in Ub pSer65 that follow the same pattern/direction as the maximum O/A dose control wells, with a higher on-target score relating to a greater phenotypic shift towards maximum O/A effect. Off-target phenotypic shift represents unexpected differences in Ub pSer65 detection, such as signal from dying cells (later described in Fig. 4D).

Fig. 2B shows the on-target phenotypic shift scores for a dose-response experiment across three replicate plates, demonstrating a strong correlation between increasing O/A dose and on target shift. To assess the robustness of the assay, Z’ values were calculated using on-target phenotypic shift scores, identifying 0.5 μM O/A as the first dose with Z’ > 0.6 across all three replicate plates (Fig. 2C) and the assay window stayed consistent with further increase to O/A concentration. In line with a previous report (Birmingham, Selfors et al. 2009), positive Z’ values indicated an acceptable assay window, with values above 0.5 indicating a very good assay window. Additionally, this O/A dose range and 5-hour treatment interval did not induce any significant toxicity as indicated by cell counts (Fig. 2D).

After establishing the mitophagy activation assay, we developed a CRISPR/CAS9 gene KO workflow to investigate gene functions in human iPSC-DA neurons, focusing on mitophagy modulation. Optimised for a 384-well format, our approach used a ribonucleoprotein (RNP) complex of CAS9 and guide RNA (gRNA) nucleofected into target cells (Fig. 3A). Gene editing was introduced on day 16, when progenitors were revived from liquid nitrogen, the only suspension step in neuron differentiation, as differentiated neurons lose viability if lifted from the matrix.

**Fig. 3:**
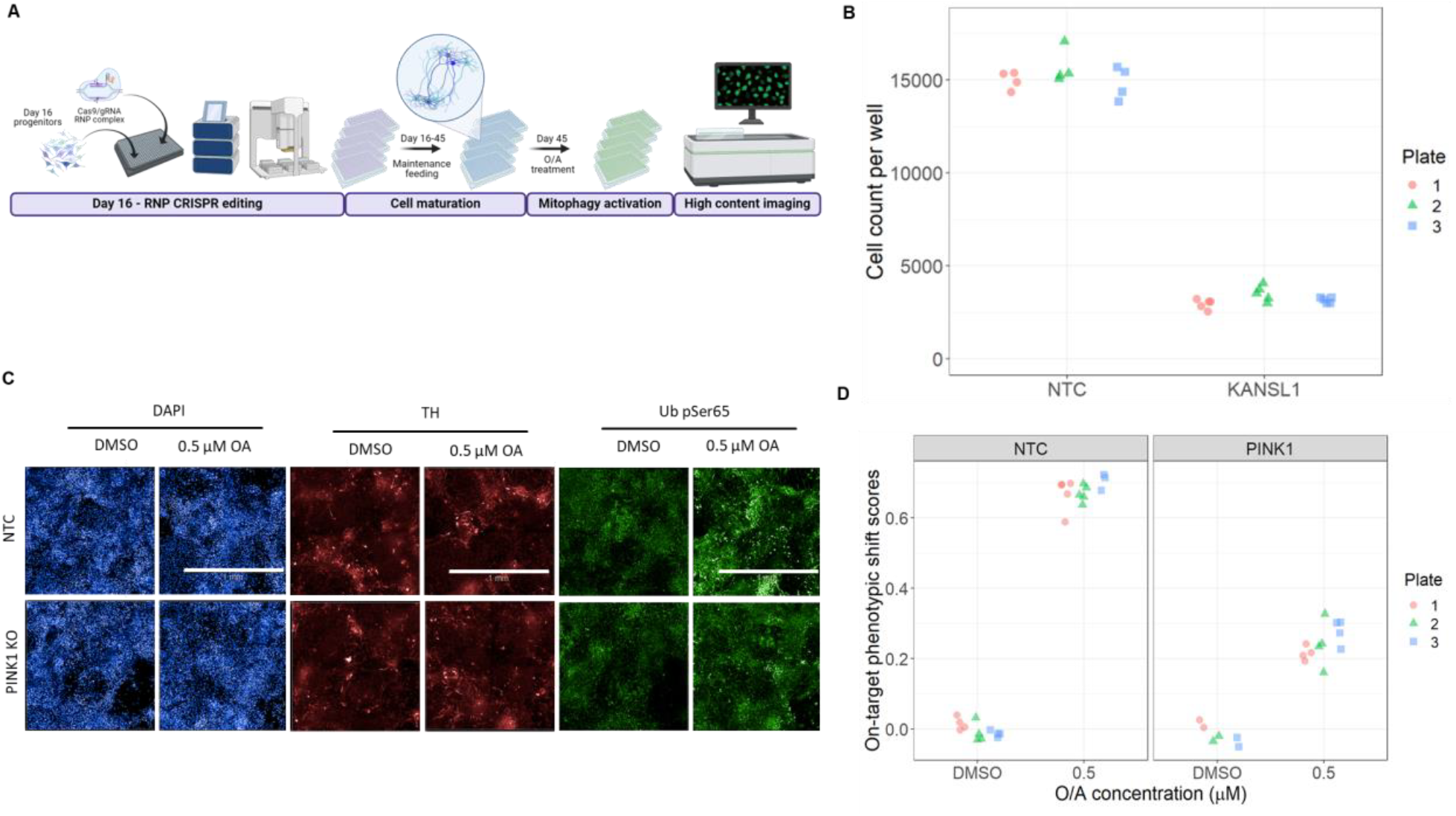
Genome editing of day 16 DA neuron progenitors using ribonucleoprotein electroporation. A) Schematic workflow of CRISPR/CAS9 gene editing in DA neuron progenitors, from day 16 RNP editing through cell maturation, mitophagy activation, and high-content imaging analysis. Created in BioRender. Carbone, B. (2025) https://BioRender.com/yupzpxi B) Cell counts per well at day 45 for KANSL1 KO and non-targeting control (NTC) across three replicate plates, demonstrating severe cell lethality in KANSL1 KO neurons. 4 replicate wells per plate for NTC control and 5 for KANSL1 KO, respectively. DMSO treated wells shown. C) Representative immunofluorescence images of non-targeting control (NTC) and PINK1 KO neurons treated with DMSO or 1mM O/A. Blue: DAPI (nuclei), Red: TH (dopaminergic marker), Green: Ub pSer65 (mitophagy marker). PINK1 KO resulted in reduced Ub pSer65 signal upon O/A treatment compared to NTC. Scale bars represent 1mm. D) Quantification of Ub pSer65 signal (on-target phenotypic shift) in response to O/A treatment across three replicate plates, as determined by AIML analysis of high-content imaging data. 2 to 5 replicates used per treatment by plate combination for each gene edit.

Gene editing was performed using a semi-automated liquid handler pipeline optimised for the generation of multiple replicate assay plates from a single master nucleofection plate – see Methods for more details.

The editing strategy relied on using three gRNAs per target, targeting a total distance of around 200bp, to maximise specificity, KO efficiency, and the likelihood of insertion-deletion mutations. Cells were maintained until day 45, when the mitophagy assay described in Fig.2 was performed. KO of KANSL1, previously reported to regulate PINK1-mediated mitophagy (Soutar, Melandri et al. 2022), was selected as a control. However, KANSL1 KO resulted in severe cell lethality, with fewer than 25% of neurons surviving to day 45 compared to the non-targeting control (NTC) gRNA (Fig. 3B). This is in line with previously reported KANSL1 phenotype (Li, Lu et al. 2022). The durable lethality of KANSL1 KO made it a suitable workflow and editing control for subsequent experiments.

Targeting PINK1, the master regulator of mitophagy (Fig. 3C) resulted in a downregulation of the Ub pSer65 signal when compared to cells transfected with NTC gRNA (Fig. 3D).

The optimised platform was then used to assess the ability of previously identified Ub pSer65 mitophagy regulators to regulate O/A-induced Ub pSer65 in human iPSC-DA neurons expressing endogenous Parkin. We selected the gene library based on a systematic review of published studies, prioritising those that explicitly reported specific modulation of UB pSer65 signal in various cell models. (Burchell, Nelson et al. 2013, Shiba-Fukushima, Ishikawa et al. 2017, Ordureau, Paulo et al. 2018, Sekine, Wang et al. 2019, Hung, Lombardo et al. 2021, Tsefou, Walker et al. 2021, Harbauer, Hees et al. 2022, Maruszczak, Jung et al. 2022, Soutar, Melandri et al. 2022, Singh, Agarwal et al. 2024). At day 45, cells were treated with 0.5μM O/A for 5 hours, prior to assessment for Ub pSer65 using high-content imaging and AIML analysis pipeline described above. Nuclear counts were used to determine whether any of target KOs induced a lethal phenotype. Severe lethality on KO of *EIF2AK1, FBOX7, KAT8, OGT*, and *ULK* was observed, with varying well-to-well variability consistently below 50% of the nuclear count when compared to the NTC gRNA control (Fig. 4B, Supplementary Fig. 2A). These genes were excluded from further Ub pSer65 modulation assessment. Fig. 4C shows representative images of Ub pSer65 signal after KO of the remaining targets in DMSO vs. O/A-treated cells. As reported previously, upon O/A treatment in NTC control cells, Ub pSer65 was upregulated, whereas this effect was reduced in *PINK1, TOM7*, and *PARKIN* KOs DA neurons. None of these gene KOs impacted DA neuron identity as assessed by TH expression (Supplementary Fig. 2B/C).

**Fig. 4:**
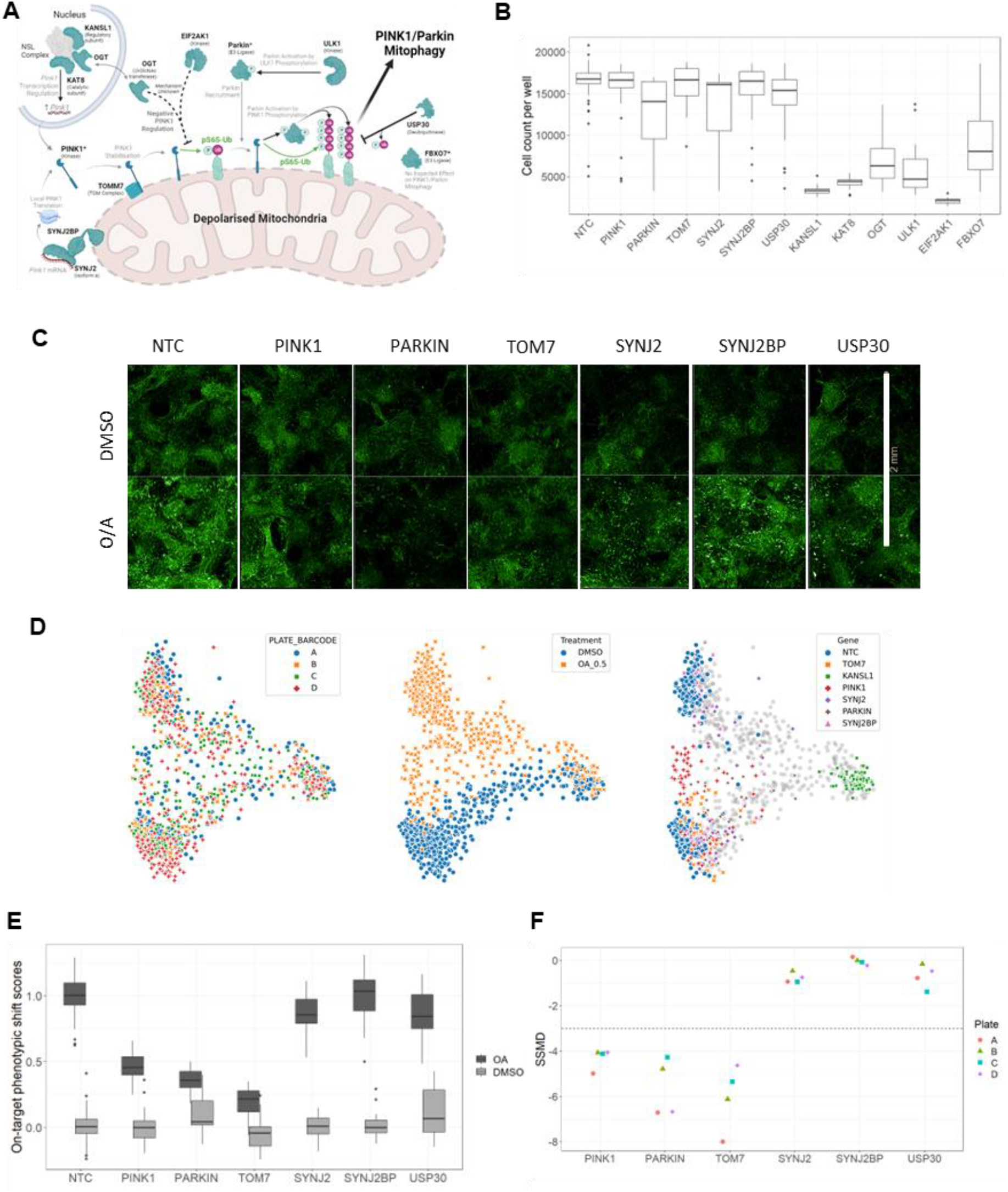
Perturbation of Ub pSer65 modulators in DA neurons. A) Schematic representation of selected targets and their reported roles in mitophagy regulation. Created in BioRender. Taylor, M. (2025) https://BioRender.com/6l91da0 B) Box plot showing cell counts per well at day 45 for each target KO and non-targeting control (NTC). Data pooled across 4 replicate plates for the visualisation, with 5 to 10 replicate wells per plate available per target KO by plate combination and 37 NTC replicates available per plate. Note that the horizontal line within each box is the median cell count per target KO. O/A treated wells shown. C) Representative immunofluorescence images of Ub pSer65 signal for selected target KOs in DMSO and 0.5 μ M O/A treated conditions. Scale bars represent 2mm. D) Principal Component Analysis (PCA) plots of machine learning features, demonstrating separation by treatment, gene condition, and replicate plate. E) Quantification of Ub pSer65 modulation (on-target phenotypic shift) for each target KO in DMSO and O/A treated conditions. Data pooled across 4 replicate plates for the visualization as in B), with both DMSO and O/A treated wells shown. F) Strictly Standardized Mean Difference (SSMD) values calculated based on on-target phenotypic shift for each target KO across 4 replicate plates, with 5 to 10 replicates available per target KO by plate combination and 37 NTC replicates available per plate and O/A treated wells used for the calculation. See Methods section for more details.

Principal Component Analysis (PCA was used to analyse and visualise machine learning-derived features (Fig. 4D). The PCA plots included all treatments and gene KOs, including lethal genes, to evaluate the performance of the model in clustering samples. The plots showed that the algorithm effectively distinguished between treatment conditions (first panel) and gene conditions (second panel), with good reproducibility across replicate plates (third panel) without plate-specific clustering. The on-target phenotypic shift for each treatment and gene KO was derived from AIML features projected along the axis representing the phenotypic shift from NTC gRNA cells in DMSO to NTC gRNA cells treated with O/A, which quantified the intended response. This projection was referred to as the ‘on-target’ score, with a score of 0 assigned to controls (NTC gRNA cells treated with DMSO) and a score of 1 to NTC gRNA cells treated with 0.5 μM O/A. The “off-target” score was derived from the perpendicular axis (rejection component), capturing morphological and signal shifts that did not correspond to the expected phenotype (E.g., *KANSL1* KO lethality). We used the on-target scores to assess Ub pSer65 modulation upon target KO (Fig. 4E). Values represented and stated below were the median phenotypic shift scores for the O/A treated wells across 4 replicate assay plates.

*PINK1* KO showed a median on-target score of 0.45 for Ub pSer65 accumulation upon treatment, confirming that PINK1 deficiency inhibited Ub pSer65 accumulation. *PARKIN* KO demonstrated a stronger response with a median on-target score of 0.36, suggesting *PARKIN* KO also impaired Ub pSer65 accumulation. This is consistent with the role of PARKIN in enhancing polyubiquitin chain formation when activated by PINK1, hence contributing additional ubiquitin substrate. *TOM7* KO showed the largest downward shift from NTC with a median on-target score of 0.22. This aligns with an established role for TOM7 in mitophagy activation (Sekine, Wang et al. 2019). As part of the TOMM complex where PINK1 accumulates during mitophagy initiation (Maruszczak, Jung et al. 2022), *TOM7* KO effectively prevented PINK1 accumulation and its role in mitophagy activation in response to O/A. In contrast, *SYNJ2* and *SYNJ2BP* KO had poor median on-target scores of 0.85 and 1.03, indicating minimal effect compared to non-targeting O/A-treated controls. Similarly, *USP30* KO yielded a median on-target score of 0.84. Although USP30 KO is known to increase Ub pSer65 levels and mitophagy (Miller and Muqit 2019, Rusilowicz-Jones, Jardine et al. 2020, Tsefou, Walker et al. 2021, Fang, Sun et al. 2023), no significant modulation of Ub pSer65 in response to *USP30* targeting was observed in the iPSC-DA neuron assay system used here.

To validate the platform for high-throughput screening, Strictly Standardised Mean Difference (SSMD) values were calculated based on on-target phenotypic shift scores (Fig. 4F). Genes were considered as hits were SSMD values were greater than 3 in absolute value (Zhang 2007). Despite some variability in cell counts and on-target scores per plate (see supplementary Fig. 3), *SYNJ2, SYNJ2BP*, and *USP30* had close to 0 SSMD values (between -2 and 2), while *PARKIN, PINK1*, and *TOM7* KOs had consistently low SSMD values (< -4), confirming the effectiveness and reliability of the platform for hit calling in high-throughput screens.

## Discussion

In this study, we established a robust, scalable platform for differentiating human iPSCs into functional midbrain iPSC-DA neurons across multiple donor lines. Characterisation at various developmental stages confirmed the efficient acquisition of key transcriptional identities and the expression of mature dopaminergic markers. Leveraging this human dopaminergic neuron model, we optimised and validated an assay system to monitor mitophagy activation through the phosphorylation dynamics of Ub pSer65, a proposed biomarker implicated in PD pathogenesis.

The generation of physiologically relevant neuronal models from iPSCs has been an area of strong interest, fuelled by the potential to recapitulate human disease biology and enable high-throughput screening efforts. However, achieving efficient and reproducible differentiation into specific neuronal subtypes has been challenging. Our optimised dual SMAD inhibition protocol, combined with precise modulation of WNT and Sonic Hedgehog signalling, yielded high percentages of midbrain DA neurons. The robust co-expression of key transcription factors (FOXA2, OTX2, LMX1) and the dopaminergic marker TH substantiated the fidelity of the differentiated neurons.

Mitochondrial dysfunction and impaired mitophagy have been implicated as key pathogenic mechanisms in PD, highlighting the importance of interrogating these processes in a disease-relevant context. Our systematic approach to optimise the detection of Ub pSer65, a specific biomarker of activated mitophagy, enabled the establishment of a robust assay window in human iPSC-DA neurons, also thanks to the machine learning model that enabled the quantification of subtle phenotypic shifts. The ability to modulate Ub pSer65 levels through pharmacological or genetic perturbations unlocks opportunities for large-scale screening efforts to identify novel regulators and potential therapeutic candidates targeting this pathway.

PINK1 and PARKIN, master regulators of mitophagy, have been studied extensively across various cellular models, including fibroblasts, HeLa cells, and iPSC-derived neurons, yet their precise roles in human iPSC-DA neurons remains relatively underexplored (Narendra, Tanaka et al. 2008, Narendra, Jin et al. 2010, Fiesel, Ando et al. 2015, Ordureau, Paulo et al. 2018, Soutar, Kempthorne et al. 2018). In this study, we combined our hiPSC-DA neuron model with a an optimised mitophagy assay. The specificity and sensitivity of the platform was validated through CRISPR/CAS9-mediated perturbation of known Ub pSer65 modulators (*PINK1, PARKIN and TOM7*) (Shiba-Fukushima, Ishikawa et al. 2017, Ordureau, Paulo et al. 2018, Sekine, Wang et al. 2019, Maruszczak, Jung et al. 2022).Ub pSer65 levels decreased upon *PINK1* and *PARKIN* KO, validating their essential roles in mitophagy. TOM7, a component of the translocase of the outer mitochondrial membrane (TOMM) complex, is required for the proper accumulation and stability of Ub at the mitochondrial membrane (Sekine, Wang et al. 2019). Consistent with previous reports, *TOM7* KO resulted in a strong reduction in Ub pSer65 levels, reinforcing its role as a key upstream regulator of PINK1-mediated mitophagy. The successful validation of *TOM7*, alongside *PINK1* and *PARKIN*, underscores the robustness and specificity of the platform in a physiologically relevant midbrain DAN context expressing endogenously expressed Parkin levels. As such, this assay platform offers a significant advancement over conventional immortalised cancer cell line models such as SHSY5Y neuroblastoma cells, which often require Parkin overexpression to observe mitophagy activation. The hiPSC-DA neurons endogenously express both PINK1 and PARKIN, providing a more accurate reflection of mitochondrial quality control processes in PD-relevant neurons. The successful adaptation of the screening platform for high-throughput genetic perturbations further reinforces its potential for systematically identifying novel regulators of mitophagy and disease-modifying targets. Additionally, these findings highlight the feasibility of leveraging this model for therapeutic screening, as the ability to modulate Ub pSer65 levels through genetic perturbations establishes a foundation for future drug discovery efforts targeting mitochondrial dysfunction in PD.

USP30 is known to play a crucial role in regulating mitophagy, acting as a deubiquitinating enzyme (DUB) that removes ubiquitin chains from mitochondrial proteins. We hypothesised that the KO of *USP30* would prevent the dismantling of ubiquitin chains, leading to an increase in ubiquitin substrate for PINK1 and elevated levels of Ub pSer65 (Tsefou, Walker et al. 2021). Although a variety of studies have shown USP30 KO/inhibition promotes mitophagy (Tsefou, Walker et al. 2021, Fang, Sun et al. 2023), we observed minimal effects on Ub pSer65 following USP30 KO. This suggests that further optimisation of the assay, such as adjusting the O/A treatment timepoints, may be needed to capture these effects.

Similarly, we investigated Synaptojanin 2 (SYNJ2) and its associated binding protein (SYNJ2BP) which regulate the transport of PINK1 mRNA to mitochondria via an RNA-binding domain in SYNJ2 (Harbauer, Hees et al. 2022, Hees, Wanderoy et al. 2024). This facilitates the transport of PINK1 mRNA from the cell soma to peripheral regions of neurons, ensuring distal mitochondria receive a continuous supply of PINK1 as needed. KO of either *SYNJ2* or *SYNJ2BP* in our DA neurons also had little effect on Ub pSer65. Recent studies have shown that AMPK activity is essential for PINK1 mRNA transport, and that insulin can inhibit AMPK to modulate this process (Soutar, Kempthorne et al. 2018, Harbauer, Hees et al. 2022, Hees, Wanderoy et al. 2024). Since B27, a common supplement used in neuronal cultures, contains high levels of insulin, this could potentially explain the lack of change in Ub pSer65 following *SYNJ2/SYNJ2BP* KO reported here.

In this study, we also identified genes that resulted in significant lethality when KO in hiPSC-DANs, namely *KANSL1, KAT8* and *EIF2AK1*.The NSL complex, involving KANSL1 and KAT8, plays a role in chromatin regulation by acetylating histone 4 lysine 16 (H4K16ac) (Radzisheuskaya, Shliaha et al. 2021). While we previously demonstrated that siRNA knockdown (KD) of either KAT8 or KANSL1 reduces PINK1 transcription and subsequent PINK1-dependent mitophagy initiation (Soutar, Melandri et al. 2022), KO of either gene in mice is embryonically lethal, which is consistent with our findings (Thomas, Dixon et al. 2008, Li, Lu et al. 2022). Additionally, KO of *EIF2AK1* has been reported to increase mitochondrial depolarisation-induced stabilisation of PINK1 and increased Ub pSer65 in HeLa SK-OV-3, U2OS and ARPE-19 cell lines (Singh, Agarwal et al. 2024), however, *EIF2AK1* KO mice were viable (Han, Yu et al. 2001). This suggests some potential non-cell-autonomous support *in vivo* or differences between neuronal and non-neuronal cell lines. KO of *ULK1* (Hung, Lombardo et al. 2021), *OGT* (Soutar, Melandri et al. 2022) and *FBXO7* (Kraus, Goodall et al. 2023) exhibited variable cell viability, suggesting that KO of these genes may impair hiPSC-DA neurons differentiation or maturation due to their involvement in a range of transcriptional processes.

Another consideration around lethality caused by gene KO is the timing of editing. Since RNP electroporation requires the cells to be in suspension, the cells were edited at day 16 and assayed at day 45. This extended interval between editing and analysis carriers a higher risk of cell loss or lethal phenotypes, as the target genes might be essential for differentiation or maturation at the earlier day 16 progenitor stage. Lentiviral delivery of CRISPR machinery, which allows more flexibility to editing timing thanks to the viral ability to transduce adherent cells, was limited due to difficulties in obtaining high-titre, uniform lentiviral preparations at scale from commercial suppliers required for high-throughput screening.

Although GWAS have identified gene variants associated with PD risk (Leonard 2025), the specific mechanisms linking these variants to disease remain underexplored. This platform provides a valuable opportunity to investigate the role of putative PD-related genes and their involvement in mitophagy. Its scalability and high-throughput format make it an attractive tool for screening PD-related disease pathways, such as endolysosomal/autophagy, and for using patient-derived disease models to explore the molecular genetics of PD.

## Summary

The studies reported here established an optimised iPSC-DA neurons model that is amenable to CRISPR-CAS9 editing and large-scale screening, offering a powerful platform for therapeutic discovery. The ability to systematically dissect disease pathways in a human midbrain dopaminergic context represents a critical step towards elucidating the aetiology of PD and develop tailored therapeutic interventions.

## Materials and methods

### Human iPSC cell culture

Human iPSC lines STBCi101-A and WTSIi075-A were supplied by European Bank for Induced Pluripotent Stem Cells (EBiSC) and the use of samples received Ethics Committee approval (CEI-117-2357). The EBiSC Bank acknowledges University of Newcastle Upon Tyne as the source of the human induced pluripotent cell line STBCi101-A (which was generated with support from the EBiSC project. The EBiSC has received support from the Innovative Medicines Initiative (IMI) Joint Undertaking (JU) under grant agreement n°115582 and from the IMI-2 JU under grant agreement No 821362, resources of which are composed of financial contribution from the European Union’s Seventh Framework Programme (FP7/2007-2013), European Union’s Horizon 2020 research and innovation programme and EFPIA.

The EBiSC Bank acknowledges Wellcome Trust Sanger Institute / HipSci as the source of the human induced pluripotent cell line WTSIi075-A which was generated with support from the EBiSC project. The EBiSC has received support from the Innovative Medicines Initiative (IMI) Joint Undertaking (JU) under grant agreement n°115582 and from the IMI-2 JU under grant agreement No 821362, resources of which are composed of financial contribution from the European Union’s Seventh Framework Programme (FP7/2007-2013), European Union’s Horizon 2020 research and innovation programme and EFPIA. The EBiSC Bank acknowledges Welcome Trust Sanger Institute as the source of the human induced pluripotent cell line which was generated under the Human Induced Pluripotent Stem Cell Initiative funded by a grant from the Welcome Trust and Medical Research Council, supported by the Welcome Trust and the NIHR/Welcome Trust Clinical Research Facility. Human iPSC line 6732BW-M3 was generated by Takara-Bio.

Vials were thawed and expanded for 2 weeks in Vitronectin (STEMCELL Technologies) coated flasks and mTeSR™ Plus media (STEMCELL Technologies), in humidified 37°C, 5% CO_2_ conditions. Media was replaced every other day and cells were passaged when reaching 80% confluency using ReleSR (STEMCELL Technologies). Cells were passaged at least twice before starting neuronal differentiation.

### iPSC to DA day 16 progenitor differentiation

Dopaminergic (DA) progenitors were generated through directed differentiation of human pluripotent stem cells over 16 days, following a modified version of established protocols (Nolbrant, Heuer et al. 2017). To optimise differentiation efficiency for each donor cell line, we tested the following neural patterning conditions:

1. 300ng/ml SHH + 0.65μM CHIR99021
2. 300ng/ml SHH + 0.8μM CHIR99021
3. 300ng/ml SHH + 0.95μM CHIR99021
4. 500ng/ml SHH + 0.65μM CHIR99021
5. 500ng/ml SHH + 0.8μM CHIR99021
6. 500ng/ml SHH + 0.95μM CHIR99021

### Immunostaining for day 14 progenitor cells

Day 14 progenitor cells were fixed using a two-step paraformaldehyde (PFA) protocol. Initially, half of the culture media was replaced with an equal volume of 2% PFA (diluted from 4% PFA stock in PBS) and incubated for 15 minutes at room temperature, followed by a second 15-minute fixation with undiluted 4% PFA. The cells were then washed 3 times with PBS, then blocked and permeabilised for 1.5 hour at room temperature with a solution of 0.125% Triton-X100 and 5% donkey serum in PBS. Primary antibodies diluted in an antibody buffer solution (0.1% Tween20 and 1% donkey serum in PBS), then incubated overnight at 4°C. Following three washes with wash buffer (0.1% Tween-20 in PBS), cells were incubated for 1 hour at room temperature with secondary antibodies and Hoechst 33342 (5 μg/mL) diluted in the same antibody buffer. After three final washes with wash buffer, the solution was replaced with PBS for imaging.

### Immunocytochemistry reagents

| Reagent/Resource | Supplier | Catalogue Number |
| --- | --- | --- |
| Formaldehyde 4%, Methanol-free | ThermoFisher | FB002 |
| D-PBS -Ca2+/-Mg2+ CTS | ThermoFisher | A1285601 |
| Triton-X | Merck Millipore | T8787 |
| Tween 20 | Merck Millipore | P1379 |
| Donkey serum | BioRad | C06SB |
| Anti-Tyrosine Hydroxylase | Merck Millipore | AB152 |
| Human HNF-3 beta /FoxA2 Antibody | R&D Systems | AF2400 |
| Human OTX2 Antibody | R&D Systems | AF1979 |
| Anti-LMX-1 Antibody, clone 4L12 | Merck Millipore | ZRB1373 |
| Human/Mouse Oct-3/4 Antibody | R&D Systems | AF1759 |
| Hoechst 33342, Trihydrochloride, Trihydrate - 10 mg/mL Solution in Water | ThermoFisher | H3570 |
| Donkey anti-Rabbit IgG (H+L) Highly Cross-Adsorbed Secondary Antibody, Alexa Fluor™ Plus 647 | ThermoFisher | A32795 |
| Donkey anti-Goat IgG (H+L) Highly Cross-Adsorbed Secondary Antibody, Alexa Fluor™ Plus 647 | ThermoFisher | A32849 |
| Donkey anti-Rabbit IgG (H+L) Highly Cross-Adsorbed Secondary Antibody, Alexa Fluor™ 488 | ThermoFisher | A21206 |
| Donkey anti-Mouse IgG (H+L) Highly Cross-Adsorbed Secondary Antibody, Alexa Fluor™ 488 | ThermoFisher | A21202 |
| Rabbit IgG, polyclonal - Isotype Control | Abcam | ab37415 |
| Recombinant Rabbit IgG, monoclonal [EPR25A] - Isotype Control | Abcam | ab172730 |
| Mouse IgG1 kappa Isotype Control (P3.6.2.8.1), eBioscience™ | ThermoFisher | 14-4714-82 |
| Mouse IgG2a kappa Isotype Control (eBM2a), eBioscience™ | ThermoFisher | 14-4724-82 |
| Goat IgG Isotype Control | ThermoFisher | 31245 |

### Mitophagy assay

The working O/A solution was prepared from individual stock solutions. Antimycin A (Abcam ab147494, 100mM solution in DMSO) was diluted to 16mM in DMSO (Mpbio 195055), while Oligomycin powder (ThermoFisher 13484164) was directly resuspended as 16mM in DMSO. Equal volumes of Antimycin A and Oligomycin were combined to form an 8mM O/A solution, which was aliquoted and frozen at –20 ° C. Each aliquot was designed to be single use used to avoid freeze thaw cycles. For cell treatment on day 45, a 2X solution of O/A was prepared in day 16 cell culture medium. Treatment was initiated by removing half the cell culture media from the cells and replacing it with the 2X O/A solution. Cells were incubated with the treatment for 5 hours at 37°C 5% CO_2_ before proceeding with fixation and immunostaining.

### Immunostaining for day 45 cells post O/A treatment

Cells were fixed using a modified paraformaldehyde (PFA) protocol optimised for tris-buffered saline (TBS) and end point assay termination. Half of the culture medium was replaced with an equal volume of 8% PFA in TBS to achieve a final concentration of 4% PFA, and cells were fixed for 30 minutes at room temperature. After three TBS washes, cells were simultaneously permeabilised and blocked for 1 hour at room temperature in TBS containing 1% donkey serum and 0.125% Triton-X100. Primary antibodies diluted in an antibody buffer solution (0.1% Tween20 and 1% donkey serum in TBS) and incubated overnight at 4°C. Following 3 washes (0.1% Tween20 in TBS), the cells were incubated with secondary antibodies in the same antibody buffer for 1 hour at room temperature. After 3 washes with wash buffer, the solution was replaced by TBS before imaging.

| Reagent | Supplier | Catalogue | Dilution in assay |
| --- | --- | --- | --- |
| UltraPure DNAase/RNAase free water | Thermalise | 10977035 | n/a |
| 10 x TBS | Sigma | T5912-1L sigma | 1X |
| Tween 20 (10%) | Sigma | 655206-50ML | 0.1% |
| Triton-X (10%) | Sigma | 93443 | 0.125% |
| Donkey serum | BioRad | C06SB | See text |
| Paraformaldehyde (16%) | Alfa Aesar | 43368 | 4% |
| Anti- Tyrosine Hydroxylase (LNC1) | Cell Signaling | #45648 | 1:250 |
| Anti- Phospho-Ubiquitin (Ser65) | Cell Signaling | #62802 | 1:1000 |
| Donkey anti-Rabbit IgG (H+L) Highly Cross-Adsorbed Secondary Antibody, Alexa Fluor™ 488 | ThermoFisher | A21206 | 1:500 |
| Donkey anti-Mouse IgG (H+L) Highly Cross-Adsorbed Secondary Antibody, Alexa Fluor™ Plus 647 | ThermoFisher | A32787 | 1:500 |
| DAPI Solution (1 mg/mL) | Thermofisher | 62248 | 1:1000 |
| Alexa Fluor™ 594 Phalloidin | Thermofisher | A12381 | 1:500 |
| IgG1 mouse monoclonal | R+D systems | MAB002R | Batch dependent, match primary Ab concentration |
| Recombinant Rabbit IgG, monoclonal [EPR25A] - Isotype Control | Abcam | ab172730 | Batch dependent, match primary Ab concentration |

### Ribonucleoprotein CRISPR-CAS9 electroporation editing protocol

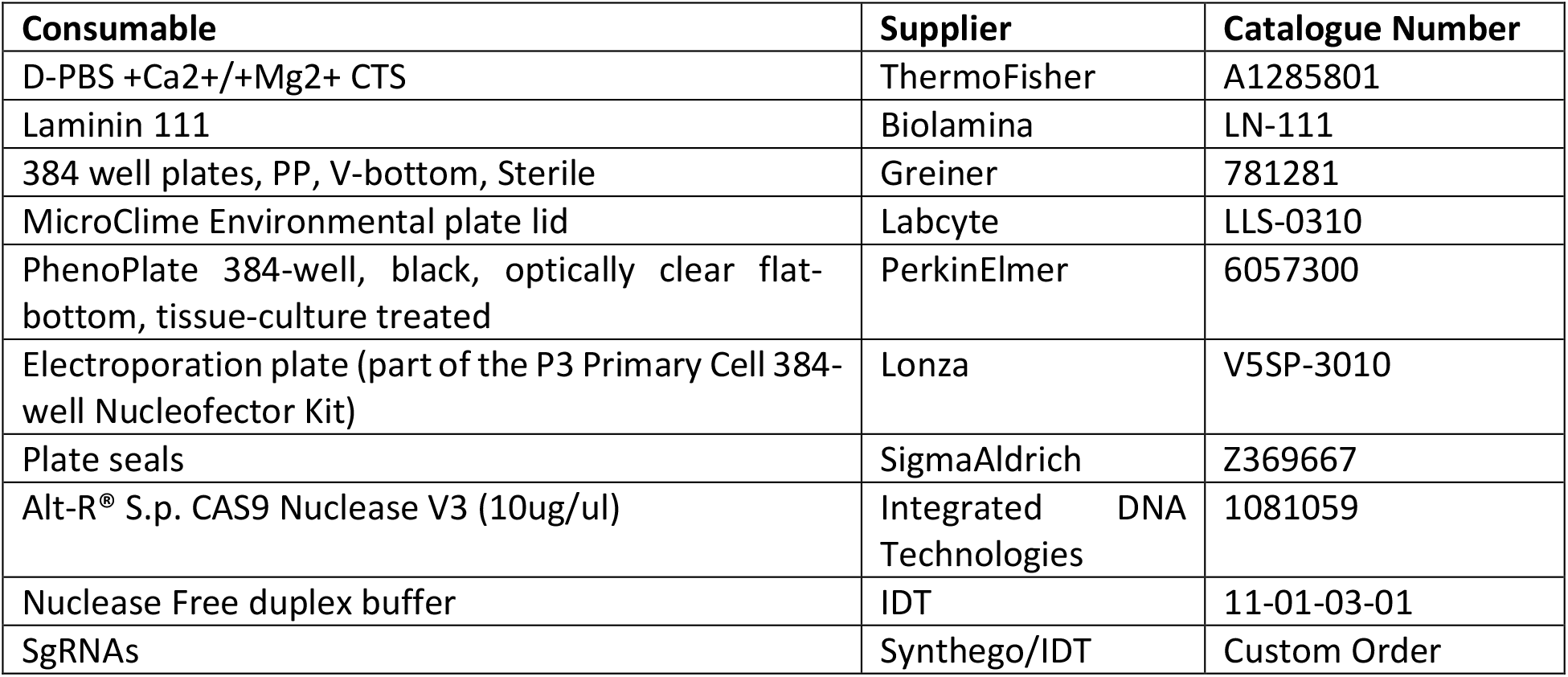

#### Day 16+ neuron culture media

Neurobasal (Thermofisher) supplemented with 1X Glutamax, 1X B27 supplement without vitamin A, Ascorbic Acid 200μM, BDNF 20ng/ml, GDNF 10ng/ml, DAPT 1μM, cAMP 0.5mM as per original protocol (Nolbrant, Heuer et al. 2017).

#### RNP workflow

384-well Phenotplates were coated with 20 μg/ml LN111 diluted in cold PBS Ca^2+^/Mg^2+^ by dispensing 27μl per 384 well and then incubated at 4°C overnight.

On the day of DA progenitors cell thaw, plates were incubated for 2 hours at 37 ° C and then washed twice with day 16 media supplemented with 10μM Rock-I.

gRNAs were resuspended in Duplex Buffer to a final concentration of 100nM by incubating overnight at 4 °C and then mixing 5 times by pipetting.

The ribonucleoprotein (RNP) complex was formed by mixing CAS9 and gRNAs in a 1:3 molar ratio and incubating the solution for 20 minutes at room temperature.

Day 16 DA progenitors were thawed in a 37 °C water bath, resuspended in day 16 media supplemented with 10μM Rock-I and counted. The cells were then centrifuged for 7 minutes at 400 g and resuspended in P3 buffer, then dispensed on top of the RNP complex. The cells and RNP solution were then transferred to a nucleofection plate for electroporation. Each electroporation reaction contained 250,000 cells and 10μg CAS9 in 20μl final volume. Cells were electroporated in a Lonza 384 high throughput nucleofector using program CL133-AA. Post electroporation, 30μl of day 16 media supplemented with 10μM Rock-I was added directly into the nucleofection plate.

The cell suspension was dispensed into the coated and pre-washed assay plates to a final concentration of 25,000 cells per well. Plates were equilibrated at room temperature for 10 minutes, then centrifuged for 20 seconds at 250 x g. Microclime lids (filled with sterile water) were used to reduce evaporation. Cells were incubated at 37 ° C, 5% CO_2_ conditions until assay day and fed every 2-3 days by performing half-media changes using day 16 media without Rock-I.

#### AIML image training and analysis

All acquired images are first pre-processed by (1) shading correction and (2) creating cell-centred patches. The cell-centred patches are created by identifying and cropping around nuclei centres from the corrected images, producing a 128 × 128 tile for each nuclei. Nuclei are identified from the Opera Phenix Harmony software using DAPI intensity, circularity and area.

A deep convolutional neural network backbone is pre-trained on the Ub pSer65 channel imaging to discriminate genetic perturbations from the processed images. ResNet18 backbone (He, Zhang et al. 2016) was used with three modifications: (i) Average Pooling and Flatten operations was replaced with a single Global Average Pooling layer to support arbitrary input image size, (ii) two dense layers of sizes 1024 and 128 are added following Global Average Pooling, and (iii) additional Mean Aggregation layer was added to enable multiple instance learning.

Following Schroff et al., (Schroff, Kalenichenko et al. 2015) we use semi-hard online triplet mining and define positive samples as wells that share same genetic perturbation and treatment condition. During training, each batch consists of K = 240 wells uniformly distributed among 12 randomly sampled genetic perturbations and stimulation conditions. For each well we further randomly sample N = 12 instances for multiple instance learning, resulting in an effective batch size of 2880.

The output of the pre-trained model is the image-based feature embedding that can differentiate between perturbations and functional states of cell populations and captures rich information on cell function and state. The phenotypic shift of each treatment and gene condition is then quantified by projecting the corresponding imaging features into the direction of NTC cells with DMSO toward NTC cells with O/A treatment. The projection yields two components: the “on-target” score and “off-target” score. The “on-target” is the projection value that is employed for the Strictly Standardized Mean Difference (SSMD) calculation.

#### Statistical methods

##### Signal window

Z-factor (Birmingham, Selfors et al. 2009), also referred to as Z’ or z-prime, is a common data quality metric for screens. Even though this metric is typically used for assay optimization, in our O/A dose-response study it is used to identify an optimal O/A concentration resulting in a good assay window, using “on-target” *phenotypic shift scores* for the calculation. The formula for the Z’ calculation is *Z*· = 1 − (3*σ*_1_ + 3*σ*_2_)/|*μ*_1_ − *μ*_2_|, where μ indicates robust mean, σ indicates robust standard deviation, and the two groups to compare could be negative and positive control, in our case the two groups are split into wells treated with DMSO vs wells treated with a particular O/A concentration. The Z’ calculation is repeated for all seven O/A concentrations tested, all wells being NTC. The calculation further uses robust summaries, so that the Z’ metric is less sensitive to outliers. It is calculated per plate out of three plates and for seven O/A concentrations, using 3 to 5 replicates per treatment by plate combination. A positive Z’ value indicates an acceptable assay window, with values above 0.5 considered desirable and indicating a very good assay window.

##### Randomisation scheme for pilot plates

DMSO and O/A treatments are to be applied in alternating columns, leaving the outer edge of the 384-well plates empty to guard against edge effects. Replicates of the target KOs were randomised to columns and vertical position (top-bottom half) of the plate to ensure good horizontal and vertical coverage across the plate.

##### Hit selection

The “on-target” projections, referred to as “on-target” *phenotypic shift scores* are the input to use for the target KO qualification as a hit. Only “on-target” *scores* for O/A treated wells are used and SSMD is calculated at the target level per plate to compare each target KO O/A treated “on-target” *score* to NTC O/A treated “on-target” scores. The formula for the SSMD calculation 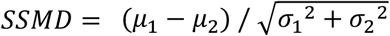 uses the robust mean *μ*_2_ and standard deviation *σ*_2_ of the NTC O/A wells per plate and similarly *μ*_1_ and *σ*_1_ of a target KO (for example PINK1) O/A wells, respectively. The SSMD calculation is performed across 12 gene edits, 6 shown in the results. Similarly to Z’ calculations, the SSMD formula utilises robust summaries to guard against outlier effects and uses a grouped version of the formula instead of replicate level calculations (Zhang 2007). 5 to 10 replicates available per target KO by plate combination and 37 NTC replicates available per plate, O/A wells used for the calculation. SSMD values greater than 3 in absolute value are considered suggesting a gene edit as a hit, i.e. significant change in Ub pSer65 signal is demonstrated if SSMD values are greater than 3 in absolute value, and not significant if SSMD values are between -3 and 3, respectively. Consistency across all four plates is desirable for hit selection.

All statistical analyses were performed in R, version 4.1.1 (R Core Team).

