## Supplementary Figures for "A CRISPR-CAS9 high throughput machine-learning platform for modulation of genes involved in Parkinson’s disease-associated PINK1-mitophagy in iPSC-derived dopaminergic neurons"

### Title

### Supplementary Figures

#### Supplementary figure 1

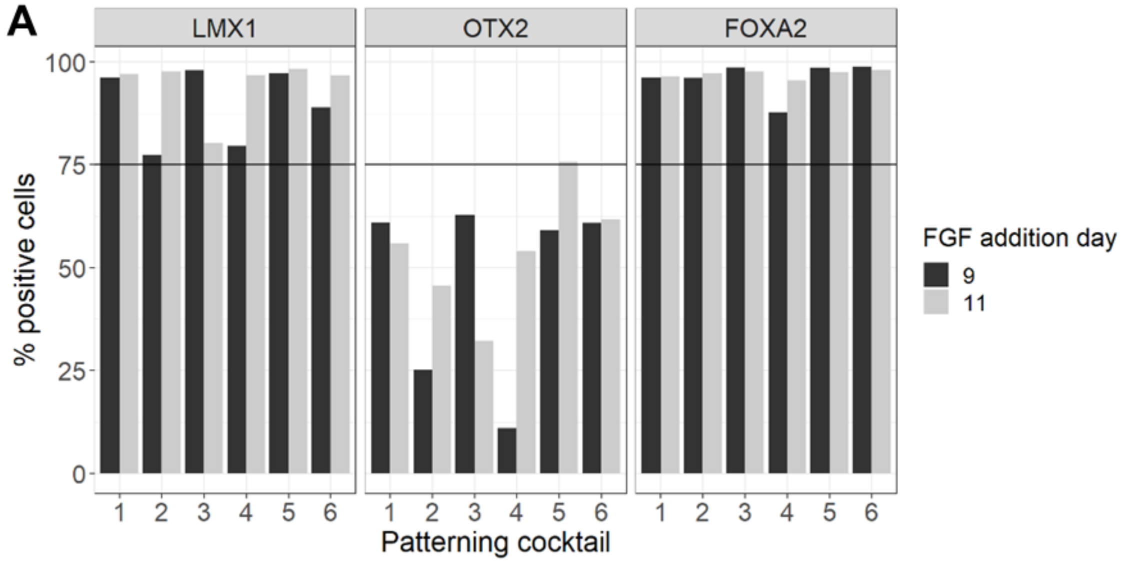

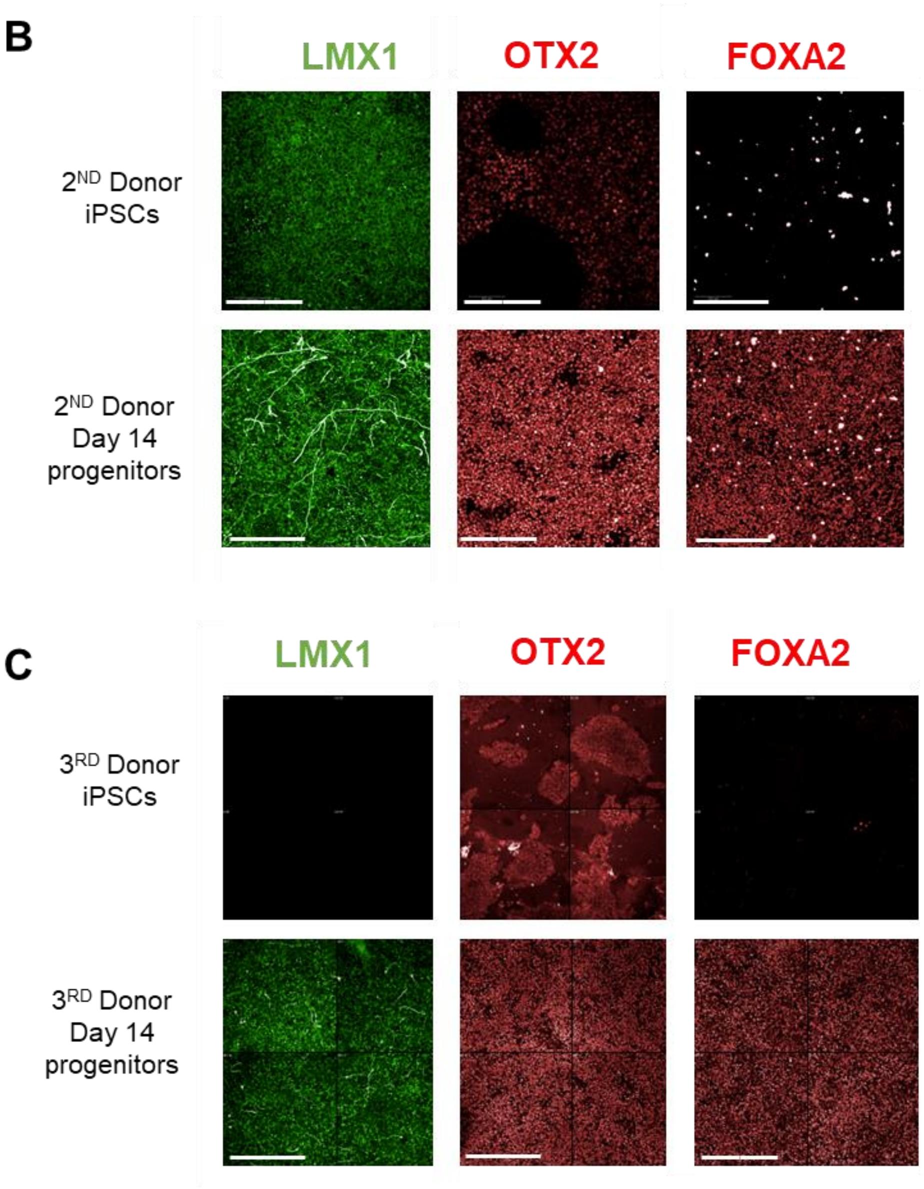


**Supplementary Fig. 1: Donor optimisation for iPSC-DA neurons differentiation**

A) Quantification of FOXA2, LMX1, and OTX2 positive cells under different patterning conditions and FGF addition times. Six different patterning cocktails were tested, with FGF added on either day 9 or day 11. The mean percentage of positive cells for each marker is shown, with 2 technical replicate wells available for FOXA2 and OTX2, and 5 for LMX1, respectively. Cocktail 5 with FGF addition at day 11 was selected as the best condition for differentiation of this donor (STBCi101-A) as it showed the highest proportion of cells expressing all three markers.

B, C) Representative images of hiPSC from two additional healthy donors (WTSIi075-A, 6732BW-M3) after differentiation towards floor plate midbrain using cocktail number 3. Scale bars represent 200µm.

### Supplementary figure 2

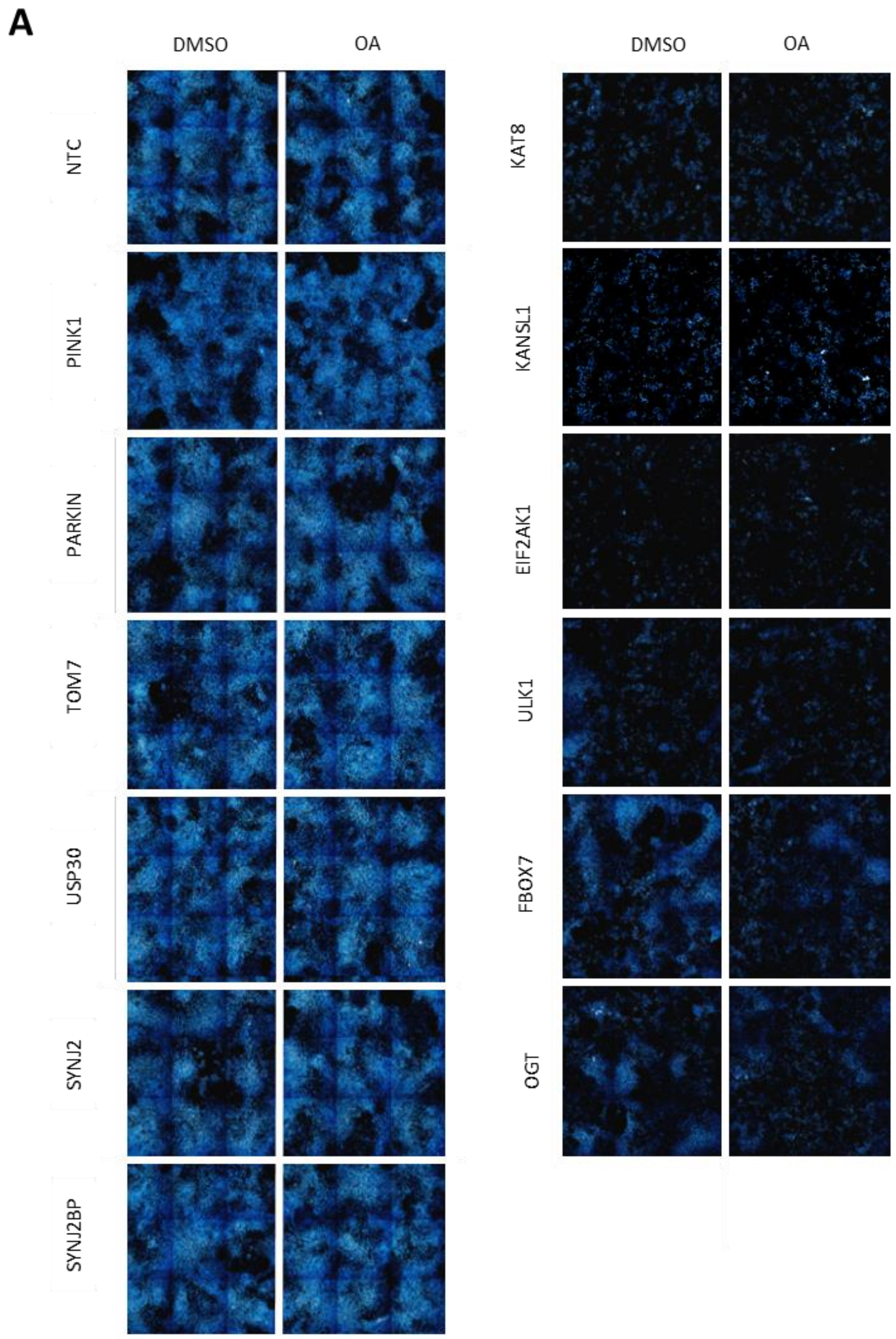

**B**

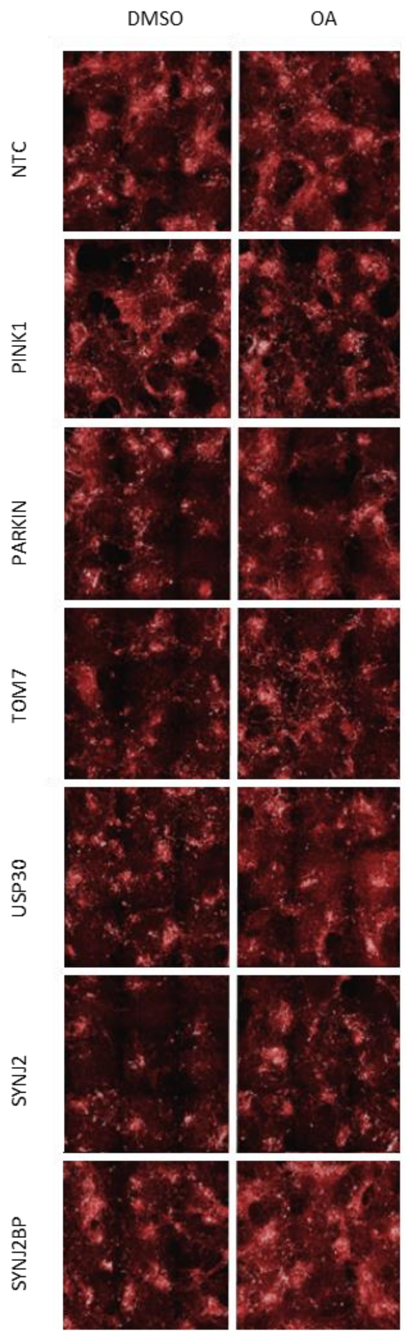

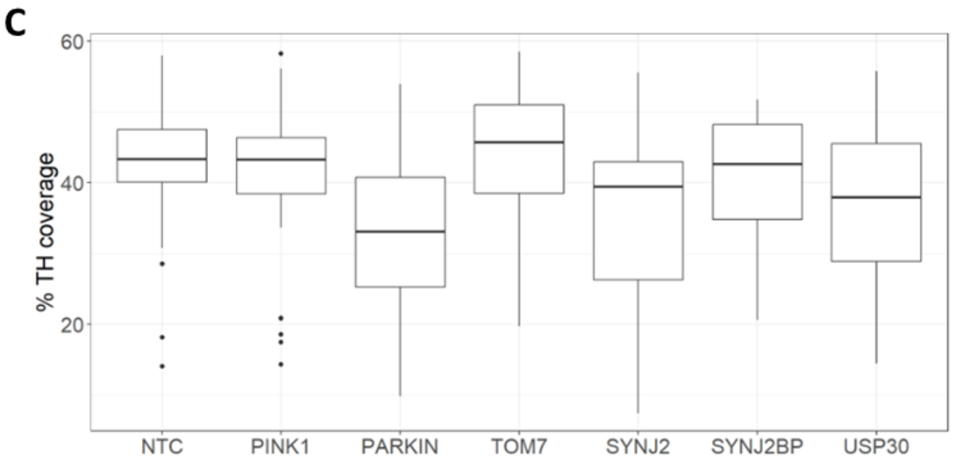


**Supplementary Fig. 2: Additional characterisation of gene KOs**

A) Representative immunofluorescence images of DAPI staining (nuclei) for all targeted genes in DMSO and 0.5 µM O/A conditions, highlighting the severe cell loss in some KOs.

B) Representative immunofluorescence images of TH staining (dopaminergic marker) for selected gene KOs in DMSO and 0.5µM O/A conditions.

C) Quantification of TH-positive cells across different gene KOs, demonstrating that KOs did not significantly impact dopaminergic neuron identity. Data pooled across 4 replicate plates for the visualisation, with 5 to 10 replicates available per target KO by plate combination and 37 NTC replicates available per plate and O/A-treated wells shown.

**Supplementary figure 3**

A

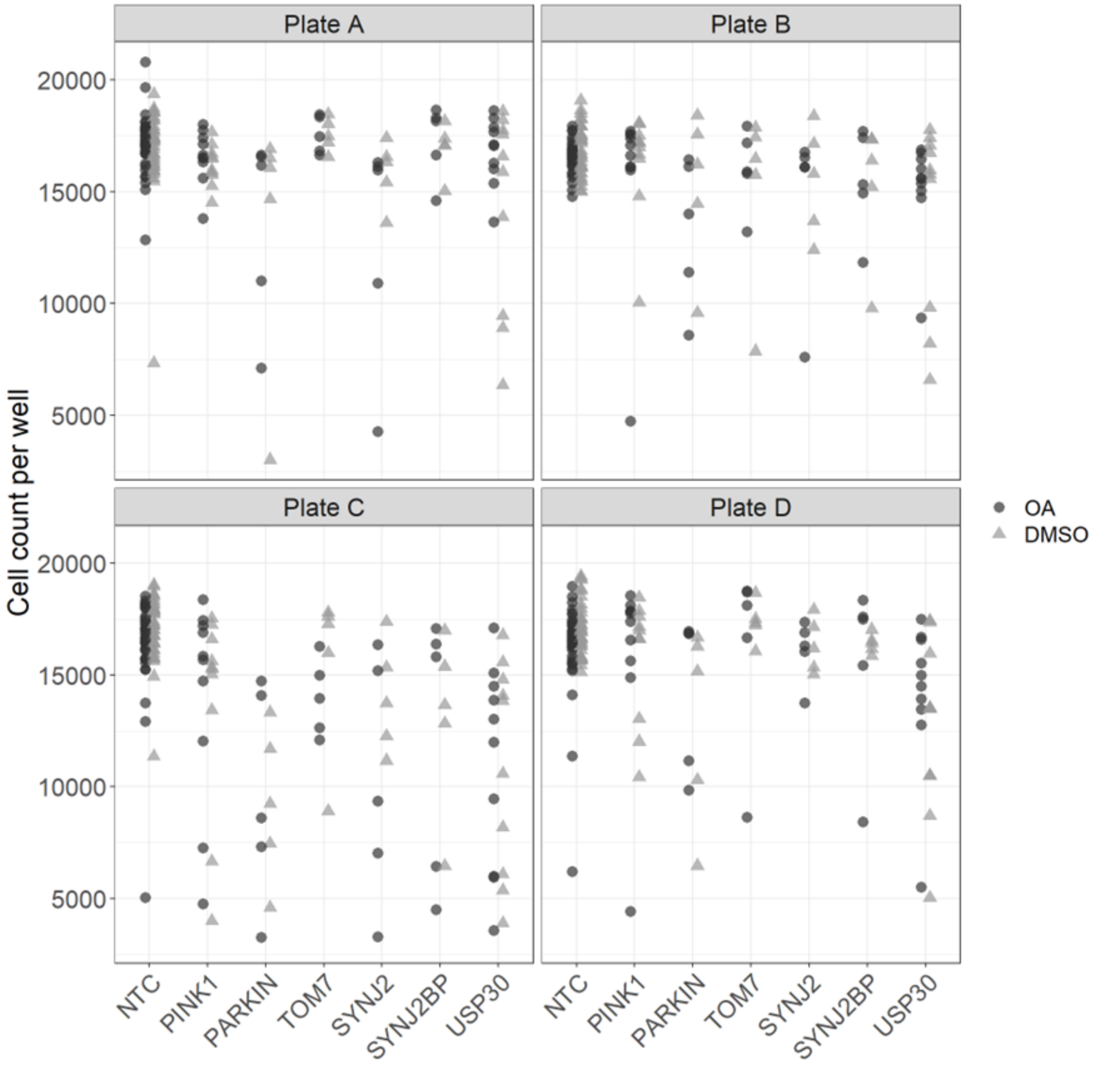

B

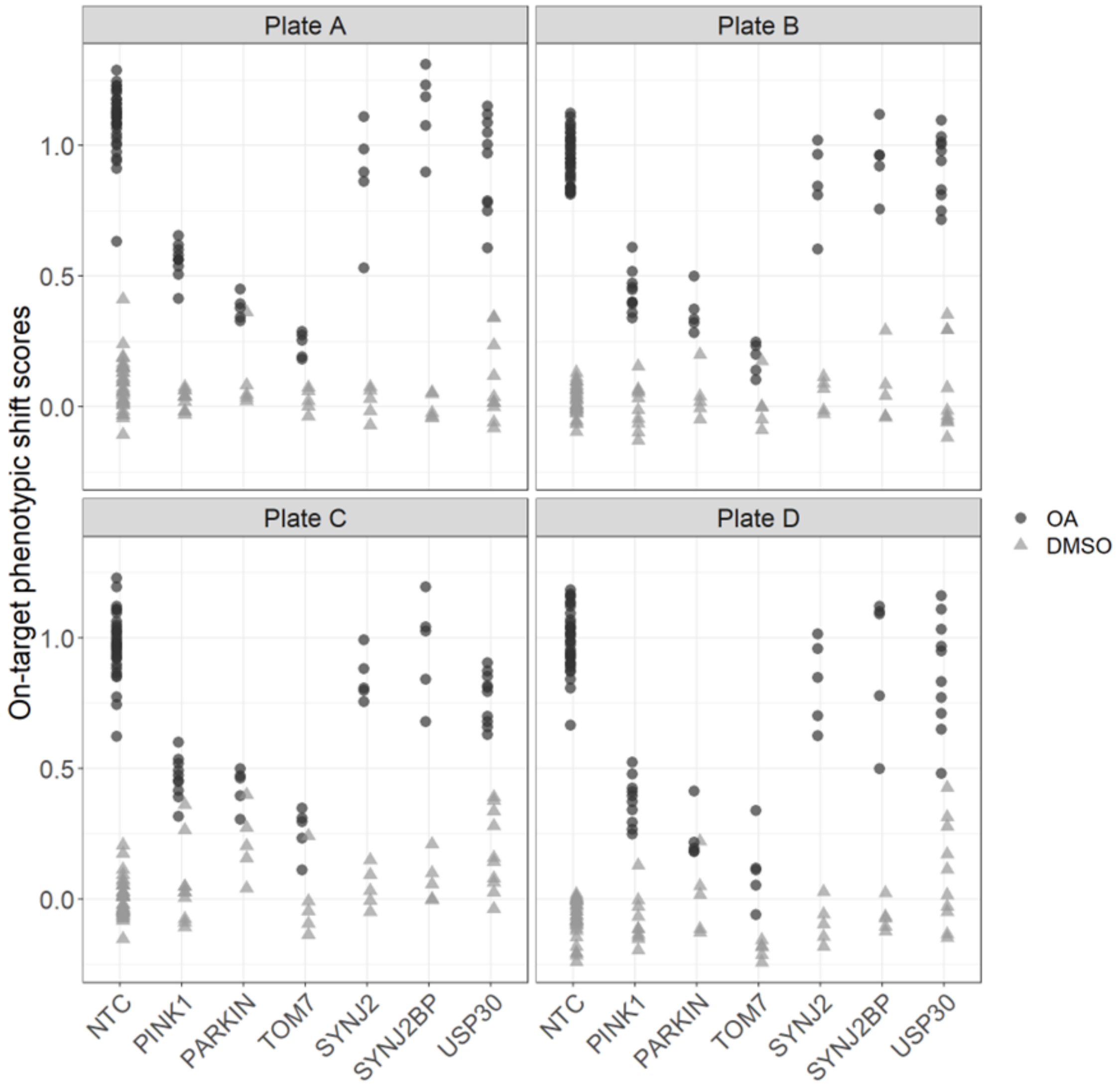


**Supplementary Fig. 3: Plate-wise analysis of gene KO effects shows minimal plate to plate variability**

A) Cell counts per well (DAPI) for each gene KO across four replicate plates (A-D) in DMSO and O/A treatment conditions.

B) On-target phenotypic shifts for each gene KO across four replicate plates (A-D) in DMSO and O/A treatment conditions.

A-B) Data across 4 replicate plates, with 5 to 10 replicates available per target KO by plate combination and 37 NTC replicates available per plate.
